# ScRNA-Seq study of neutrophils reveals vast heterogeneity and breadth of inflammatory responses in severe COVID-19 patients

**DOI:** 10.1101/2021.12.01.470817

**Authors:** Jintao Xu, Bing He, Kyle Carver, Debora Vanheyningen, Brian Parkin, Lana X. Garmire, Michal A. Olszewski, Jane C. Deng

## Abstract

Severe cases of COVID-19 are characterized by dysregulated immune responses, but specific mechanisms contributing to the most severe outcomes remain unclear. Neutrophils are the most abundant leukocyte population in human hosts and reach markedly high numbers during severe COVID-19. However, a detailed examination of their responses has been largely overlooked in the COVID-19 literature to date. Here, we report for the first time a dedicated study of neutrophil responses using single-cell RNA sequencing (scRNA-Seq) of fresh leukocytes from 11 hospitalized adult patients with mild and severe COVID-19 disease and 5 healthy controls. We observed that neutrophils display a pronounced inflammatory profile, with dramatic disruption of predicted cell-cell interactions as the severity of the disease increases. We also identified unique mature and immature neutrophil subpopulations based on transcriptomic profiling, including an antiviral phenotype, and changes in the proportion of each population linked to the severity of the disease. Finally, pathway analysis revealed increased markers of oxidative phosphorylation and ribosomal genes, along with downregulation of many antiviral and host defense pathway genes during severe disease compared to mild infections. Collectively, our findings indicate that neutrophils are capable of mounting effective antiviral defenses but adopt a form of immune dysregulation characterized by excess cellular stress, thereby contributing to the pathogenesis of severe COVID-19.

## Introduction

Severe lung injury and systemic inflammation are the main hallmarks of severe respiratory viral infections, including SARS-CoV-2 (1). Neutrophils, or polymorphonuclear leukocytes (PMNs), are the most abundant leukocyte population in the blood and are found in high numbers in the lung during severe respiratory viral infections (2). Although neutrophils are considered as a primary driver of the pathogenesis of ARDS (acute respiratory distress syndrome) and have been implicated in the pathophysiology of severe COVID-19 (3–7), the challenges of isolating PMNs for analysis and their inability to survive cryopreservation have resulted in a poor understanding of genomic and phenotypic heterogeneity of neutrophils during COVID-19 infection and disease progression (8). While recent studies have started to acknowledge the heterogeneity of neutrophils during cancer and other chronic diseases (9, 10), neutrophils are still largely assumed to be a homogenous cell population with a limited armamentarium of effector functions, such as ROS production, neutrophil extracellular trap (NET) release, and bactericidal activity. It is our goal to investigate whether neutrophils are capable of adopting different phenotypes during acute respiratory viral infections and to identify dysregulated immune pathways that contribute to the persistent immunopathology underlying severe COVID-19 infections.

Single-cell RNA sequencing (scRNA-seq) analysis of the peripheral immune response to SARS-CoV-2 has provided novel insights into immune cell heterogeneity and dysregulation during COVID-19 (5, 11–17), including identifying enrichment in classically activated monocytes and dysregulated T cell responses (18, 19). However, most of those studies have focused on the analysis of preserved peripheral blood mononuclear cells (PBMC), which are mainly comprised of monocyte and lymphocyte populations. Only a few studies reported an incomplete and possibly biased picture of increased immature and dysfunctional neutrophils in the PBMC fraction, identified as low-density neutrophils (LDNs) (12, 20, 21). Thus, it remains unclear whether distinct neutrophil populations exist, and if so, how they change for COVID-19 progression or with disease severity.

To fill this important void in the knowledge of neutrophil biology, we analyzed neutrophils from hospitalized adult patients with mild and severe COVID-19, by using an unbiased and comprehensive approach to transcriptomic analysis of unpreserved, whole peripheral blood leukocytes including neutrophils at the single-cell level. We hypothesized that this approach would reveal neutrophil subsets marked by differential antiviral and inflammatory gene expression profiles, which would be associated with the presence and severity of infection. Our analysis revealed that quantitatively and qualitatively, neutrophils could be a more robust driver of inflammatory responses than monocytes, underscoring the importance of investigating the considerable heterogeneity of responses in the neutrophil population. We also identified potential cross-regulatory mechanisms by which neutrophils may interact with other leukocytes, with a notable dropout in the number of predicted cross-regulatory pathways with increasing disease severity, suggesting loss of immune regulatory mechanisms. Collectively, our results fill an important information gap in the COVID-19 literature, which has largely focused on monocyte and lymphocyte responses. The design of immunotherapeutic approaches, therefore, should account for the contributions of neutrophils, with future studies focused on understanding the factors that regulate different aspects of neutrophil activation during severe respiratory infections.

## Results and discussion

### Neutrophils are major contributors to the inflammatory response relative to other peripheral leukocytes during COVID

Activated monocytes and T cells have been portrayed as the primary cellular drivers of inflammation during severe COVID-19. Neutrophils, despite being the predominant leukocyte population in terms of numbers (22), have been largely overlooked due to the inability of these cells to survive long-term storage and cryopreservation. Because of the paucity of neutrophil data from COVID-19 infected subjects, we performed droplet-based scRNA-seq (10X Genomics) to examine the transcriptomic profiles of peripheral neutrophils and other leukocytes from 11 hospitalized adult COVID-19 patients and 5 healthy donors (HD) (Fig. S1A&S1B). To reduce confounding, we excluded subjects with immunosuppression, autoimmune disease, chronic infection, active malignancy, or other significant comorbidities. The 11 patients with COVID-19 were classified into two groups based upon severity – “mild” (n□=□4, hospitalized but needing ≤50% O_2_), or “severe” (n□=□7, hospitalized but needing > 50% O_2_ or in Intensive Care Unit). The clinical characteristics of enrolled patients are detailed in Fig. S1B. All patients were of the same gender with no statistical differences in age between groups. PMNs and other leukocytes were isolated from peripheral blood samples, with the first collection done within 72 hours of hospital admission (“T1”) and the second 5-7 days later if the patients were still hospitalized (“T2”). Since neutrophils are particularly sensitive to degradation, isolated cells were immediately processed for scRNA-seq experiments (see Methods). Other leukocytes from peripheral blood were included at approximately equal proportions in the scRNA seq analysis to dissect cell-cell interactions.

A unified single-cell analysis pipeline was employed, including preprocessing involving batch removal and quality control steps (see Methods in Supplementary files). A total of 108,597 high-quality cells from all samples proceeded to downstream analysis. Among these cells, 30,429 cells (28%) were from the healthy donor group, 22,188 cells (20%) were from the mild group, and 55,980 cells (52%) were from the severe group. Using Seurat (23) and SingleR (24), we identified 15 major cell types or subtypes according to the expression of canonical gene transcripts in the peripheral blood (Fig. 1A&1B). Among them, 45,463 cells are classified as mature (CXCR2^+^ FCGR3B^+^) or immature (CD24^+^PGLYRP1^+^CEACAM8^+^) neutrophils (Fig. 1A&1B) (7, 18).

**Fig. 1.**
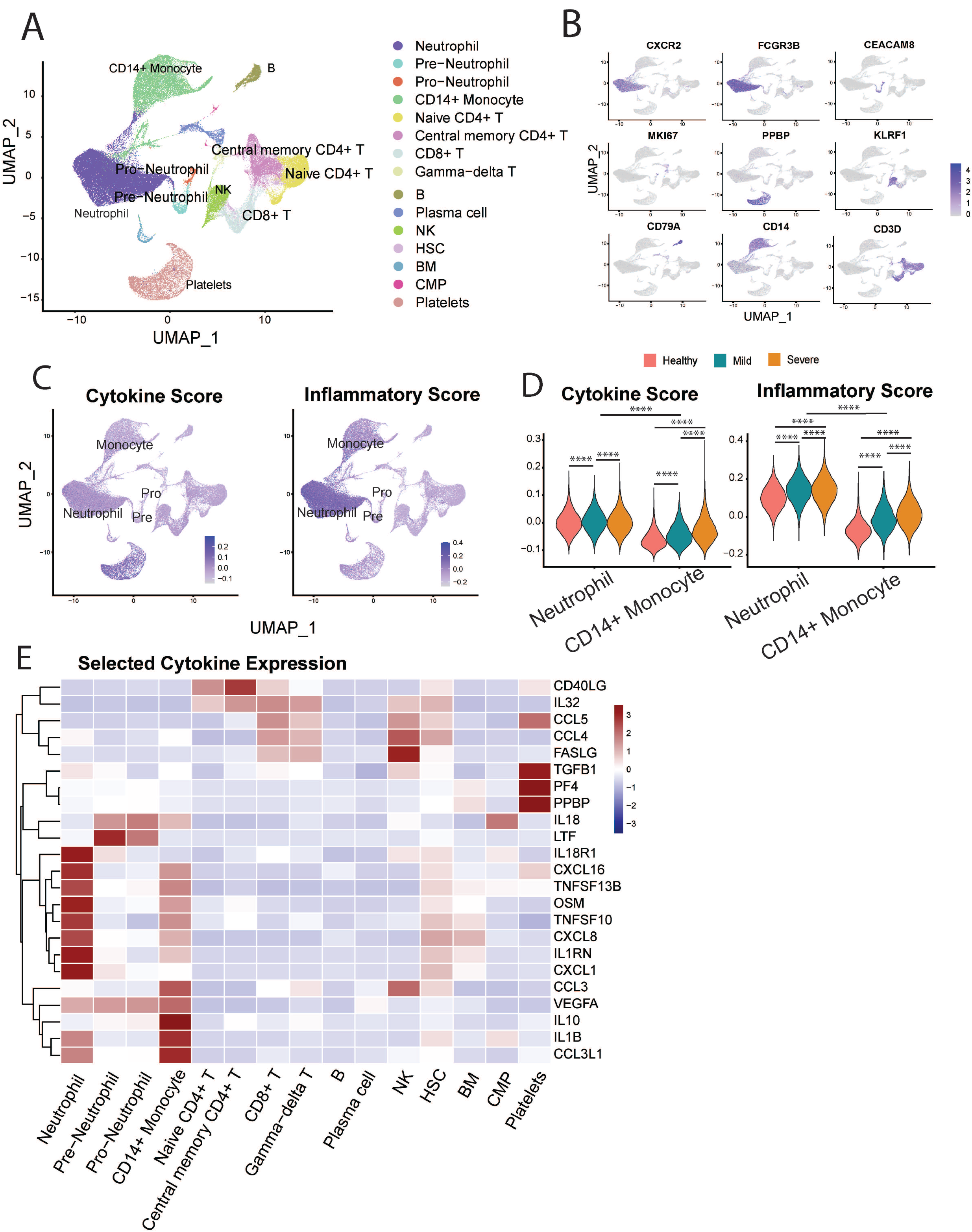
Neutrophils display marked inflammatory signatures relative to other leukocyte populations. A) Cellular populations identified by scRNA seq. The UMAP projection from HD (n□=□5), Mild (n□=□4), severe (n□=□6) samples. B) canonical cell-defining transcripts were used to label clusters by cell identity as represented in the UMAP plot. Data are colored according to expression levels and the legend is labeled in log scale. C) UMAP plots of cells colored by cytokine score (left) and inflammatory score (right panel). D) Violin plots indicate progression of cytokine (left) and inflammatory scores (right panel) for neutrophils and monocytes with increased severity of infection. E) Heatmap of selected cytokine genes in different subsets of cells. *p < 0.05, ** < 0.01, *** < 0.001.

Next, we analyzed the transcriptional profiles of myeloid (monocyte and neutrophil) populations to determine the differential contributions of each cell type towards the inflammatory landscape during COVID-19. First, to validate the compatibility of our approach with previous studies, we examined transcription of two monocyte factors most consistently reported to change with COVID-19 severity. Monocytes from severe COVID-19 patients downregulated HLA-DRA and upregulated CD163 (Fig.S2A&B), consistent with previous reports (12, 17). Next, we defined cytokine and inflammatory pathways scores based on the overall expression of cytokine and inflammatory response genes (Figure 1C and 1D, Supplementary Table 1) (17, 25). Monocytes and megakaryocytes are considered to be major sources of systemic cytokines production (17). Surprisingly, we observed that neutrophils have greater potential to regulate the magnitude of the systemic inflammatory response, indicated by their high cytokine and inflammatory pathway scores (Fig. 1C-1D). In addition, neutrophils outnumber monocytes by 10 to 40-fold (Fig.S2C). Altogether, the greater number of neutrophils and their higher inflammatory pathway scores suggest that neutrophils are a major contributor to the magnitude and quality of systemic inflammatory responses in COVID-19.

We then investigated specific inflammatory gene signatures for each cell subtype and found neutrophils have distinct inflammatory cytokine/receptor profiles with enrichment of *CXCL1, IL1RN, CXCL8, TNFSF10, TNFSF13B, CXCL16*, and *IL8R1* (Fig. 1E). Immature neutrophils express markedly higher levels of lactoferrin (*Ltf*) and *Il18* (Fig. 1E). Furthermore, we found strong transcriptional upregulation of *S100A9* and *S100A8* alarmins in neutrophils from COVID-19 patients (Fig. S2D), previously reported to correlate with disease severity (13, 26). This increase persists for *S100A8* in the severe compared to the mild COVID-19 group (Fig. S2E). Together, our data show that neutrophils can be major contributors to the inflammatory response relative to other peripheral leukocytes during COVID.

### Identification of dysregulated neutrophil phenotypes in severe COVID-19 patients

We next analyzed transcriptional changes within the overall neutrophil population associated with the severity of the disease. Neutrophils from healthy, mild, and severe patient groups show distinct differentially expressed genes (Fig. 2A), reflecting significant transcriptomic changes during disease progression. Neutrophil transcripts which are robustly expressed in the uninfected state, including anti-proliferation and pro-apoptotic genes (*LST1, G0S2, CPPED1, BTG2, PMAIP1*) and anti-inflammatory genes (*AMPD2, SEC14L1, ZFP36*), are significantly downregulated in COVID-19 patients (Fig 2B). Neutrophils from mild patients have increased expression of genes associated with anti-viral responses, including Interferon stimulated genes (ISGs) and TRAIL (*TNFSF10*) (Fig. 2B). Remarkably, expression of these genes is attenuated in PMNs from the severe patients, whose neutrophils displayed increased activation markers including *GBP5* (activator of NLRP3 inflammasome assembly) (27) and *CD177*, previously associated with COVID-19 severity and death (28), as well as stress-related genes such as IRAK3, FKBP5, IL1R2 (Fig. 2B).

**Fig. 2.**
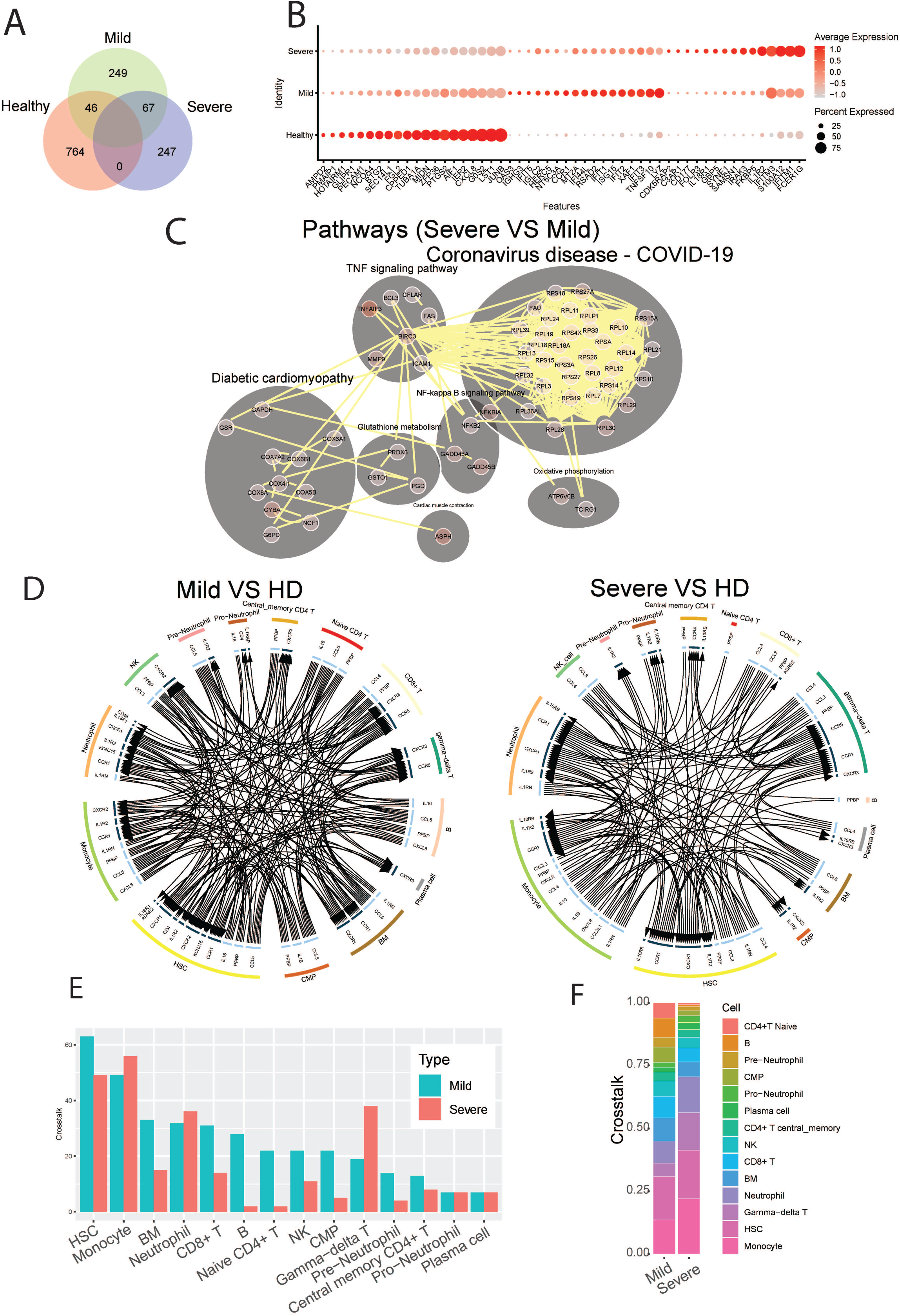
Identification of dysregulated neutrophil phenotypes in severe COVID-19 patients. A) Venn plot of significantly up-regulated genes (adjusted P-value <0.05) in neutrophils from healthy controls, mild and severe COVID19 patients. B) Top 10 differentially expressed upregulated genes in neutrophils from healthy controls, mild and severe COVID19 patients, respectively. C) Predicted cell-cell interaction networks of significantly up-regulated pathways (adjusted P-value <0.05) in neutrophils from severe COVID19 patients compared with that from the mild group. D) Circos plot showing the up-regulated cellular crosstalk mediated by significantly (adjusted P-value <0.05) up-regulated ligand-receptor pairs between inflammation-related cell types from mild or severe COVID19 patients compared with that from healthy controls. E) Count of the up-regulated cellular crosstalk for every cell type in mild and severe COVID19 patients, respectively. (F) Composition of the up-regulated cellular crosstalk in mild and severe COVID19 patients, respectively.

To further investigate how neutrophils may functionally differ during infection as compared to healthy controls, we performed pathway analysis of neutrophil transcriptomes. In hospitalized patients with milder COVID-19 disease, we observed broad activation of multiple pathways involved with immune responses to various viral infections, including COVID-19 related pathways (mostly antiviral genes), NOD-like receptor signaling, TLR signaling, and immune responses to both viral and intracellular pathogens (e.g., influenza A, Salmonella, Epstein-Barr) (Fig S3). Thus, the diversity of upregulated pathways highlights a more pronounced antiviral neutrophil response in hospitalized patients with milder COVID-19. Conversely, in neutrophils from patients with severe disease, we observed significant activation of NF-kB signaling, and TNF signaling pathways, as well as oxidative stress response pathways (e.g., cyclooxygenase genes, glutathione metabolism, and oxidative phosphorylation), compared to those from mild COVID-19 patients (Fig 2C). This suggests stress response phenotype in severe patients, rather than a protective-antiviral phenotype seen in the moderate disease. Notably, unlike patients with mild disease, severe patients show marked induction of ribosomal genes, suggesting an increase of cellular protein production capacity beyond the observed increase in active gene transcription.

Finally, to provide immunologic context for how neutrophils interact with other cell types, we conducted an analysis on the intercellular crosstalk between cytokines and receptors on immune cells. To identify how cytokine-receptor-mediated cell-cell interactions (CCI) evolve across disease severity, we counted the active CCIs that represent the active intercellular communication probabilities between up-regulated cytokines and receptors on all cell types in different COVID-19 statuses. We found that during mild disease, there are overall more active CCIs among all of the different cell populations than that in severe disease (Fig. 2D and E, S4). Conversely, during severe disease, many of these active CCIs drop out, resulting in potential degradation of cell-cell cross-regulatory mechanisms. Cell-cell interactions become concentrated and are dominated by interactions between 4 major cell types: neutrophils, monocytes, gamma-delta T cells, and hematopoietic stem cells (HSC), which accounts for more than 60% of the cell-cell interactions in severe disease (Fig. 2F). As the illness proceeds, we found that in mild patients who recovered from the disease, diverse cell-cell interactions remain preserved at later timepoints, while severe patients who eventually succumb have progressive loss of cell-cell interaction diversity (Fig. S4). These data support that neutrophil cell-cell interaction become progressively dysregulated in patients with severe COVID-19, and that a potential mechanism by which neutrophils contribute to severe inflammation may be a reinforcement of activation pathways for specific cell populations such as monocytes and gamma-delta T cells, which have been reported to be robustly pro-inflammatory cell populations during viral infections.

### COVID-19 resulted in alterations of neutrophil subset compositions and their transcription profiles across the levels of the disease severity

We next examined whether different phenotypes of neutrophil populations could be identified by scRNA-Seq. We performed cluster analysis of neutrophil scRNA data using the SNN modularity optimization-based clustering algorithm. In total, 9 distinct clusters of neutrophils could be identified based on specific patterns of gene expression. Cluster 9 represents pro-neutrophils (*DEFA3*^*+*^*DEFA44*^*+*^*MPO*^*+*^*ELANE*^*+*^ *AZU*^*+*^; azurophilic granule content genes), cluster 7 represents pre-neutrophils (*LTG*^*+*^*LCN2*^*+*^*CAMP*^*+*^*MMP8*^+^; specific and gelatinase granule content genes), and the remaining 7 clusters represent mature neutrophils (*CXCR2*^+^) (Fig 3A and 3B). The two immature neutrophil clusters (clusters 7 and 9) exhibit robust gene expression of their respective granule content proteins but relative suppression of all of the other genes (Fig 3C). Conversely, the mature neutrophil clusters had suppression of granule content genes, but distinct patterns of gene activation that were relatively decreased in the immature populations (Fig 3C). Clusters 2 and 8 displayed upregulation of MMP9, several S100A genes including S100A12, and *MME* (i.e., Neprilysin), all of which have been implicated in the pathogenesis of COVID-19 (29, 30). Clusters 3, 5, and 6 had high levels of expression of regulatory genes for transcription and apoptosis. Notably, cluster 4 was significantly enriched in interferon (IFN) stimulated genes (ISGs, e.g., *ISG15, IFIT* genes, *MX1, GBP*, G*BP5, HERC5*, and *RSAD2*). Thus, contrary to the assumption that they are a homogenous and transcriptionally quiescent cell population, mature neutrophils display the ability to activate differential gene expression programs, ranging from inhibitory/regulatory subsets to a preferentially antiviral subset with activated IFN-regulated gene expression program.

**Fig. 3.**
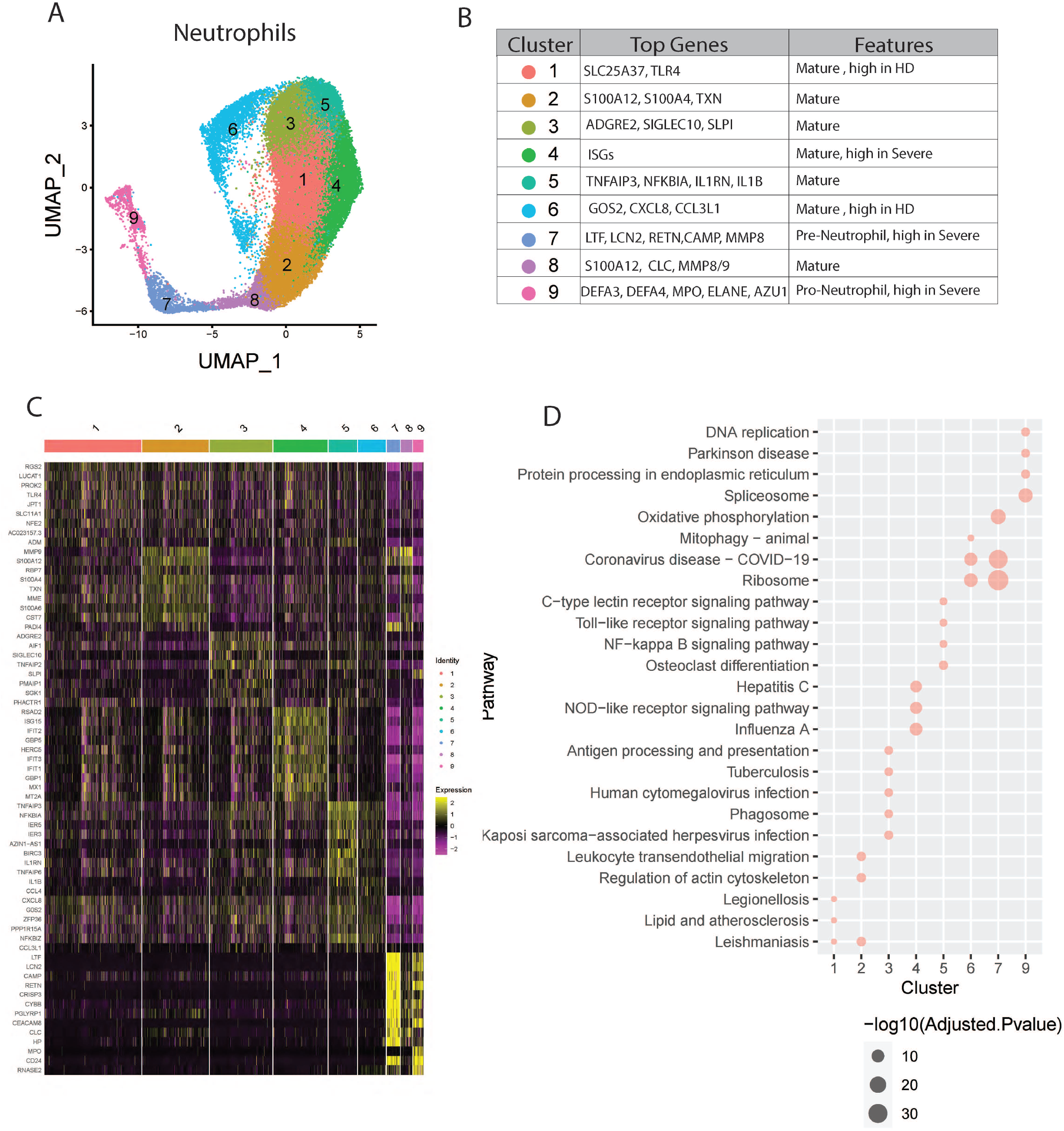
Neutrophil heterogeneity in COVID-19 patients. (A) UMAP plot of neutrophil clusters. (B) Nomenclature and marker genes for each neutrophil cluster. (C) Top5 up-regulated genes in every neutrophil cluster. (D) Top3 pathways significantly enriched in up-regulated genes in every neutrophil cluster.

Subsequent pathway analysis provided insights about possible biological functions of each neutrophil subset (Fig. 3D). Pathway analysis of cluster 4 revealed significant activation of viral response pathways as well as NOD-like receptor signaling pathway, supporting its distinct role in anti-viral responses. Other clusters also show specific pathway enrichment, for example, cluster 1 exhibited pathways involved in ferroptosis, cluster 3 and 5 in NF-kappa B signaling, and cluster 7 with activated Ribosome and Coronavirus disease-COVID-19, which is consistent with the concept of “pre-neutrophils” being robustly mobilized during acute infection with SARS-CoV-2.

Next, we determined whether all of these clusters exist at baseline and whether specific neutrophil subsets were augmented depending on the presence or severity of infection. We found higher proportions of clusters 1 and 6 in healthy compared to infected subjects, while clusters 4, 7, and 9 were increased in COVID-19 patients, especially in the severe group (Fig. 4A, Fig. S5A-B). Since clusters 7 and 9 are immature neutrophils, their increase provides evidence of emergency myelopoiesis in severe COVID-19 patients, also supported by previous reports (12, 26). Overall, cluster 4, enriched with anti-viral responses, is significantly associated with disease severity, while cluster 6 was negatively associated with the severity of the disease (Fig. 4B). Within each cluster, we also observe the abilities of neutrophils to up-or down-regulating pathways based upon disease state. For example, compared to healthy controls, cluster 7 neutrophils (immature neutrophils) from infected subjects upregulate genes involved in multiple pathways associated with host defense, including neutrophil extracellular trap formation, cytokine signaling, and NF-kB signaling. Cluster 1 and 4 PMNs from infected subjects activate COVID-19 and ribosome pathway-related genes (Fig. 4C), while subjects with severe infection have further activation of the ribosome and COVID-19 pathways, as well as oxidative phosphorylation in multiple neutrophil clusters compared to mild disease (Fig. 4D). Interestingly, neutrophils display progressively decreased activation of hepatitis, influenza, and other viral pathways with worsening COVID-19 disease severity (Fig. 4D). Strikingly, cluster 6, which is decreased in proportion during infection, displays downregulation of multiple pathways during severe disease, including those related to IL-17 signaling, NF-kB, and cAMP signaling (Fig. 4D).

**Fig. 4.**
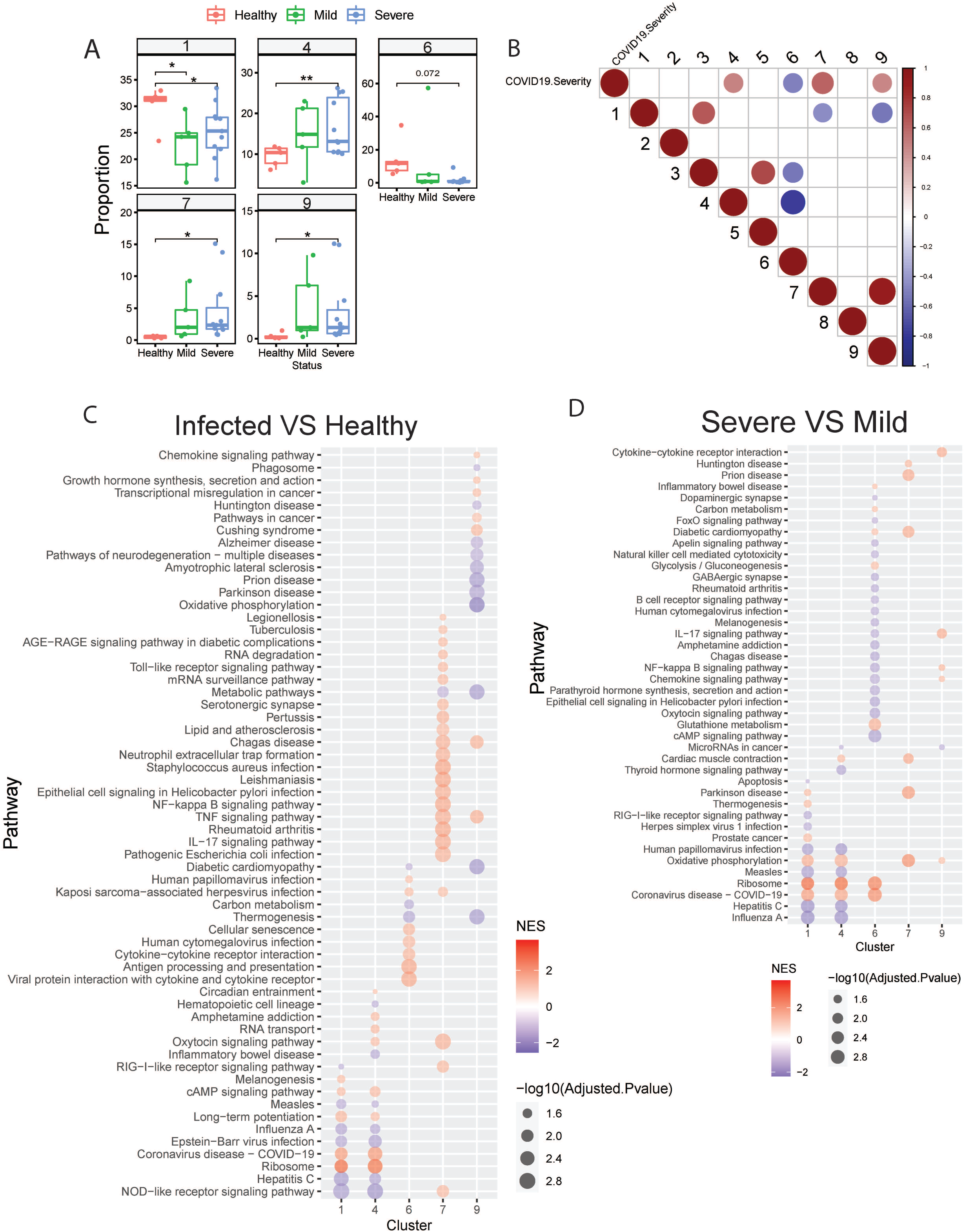
COVID-19 resulted in alterations of neutrophil subset compositions and their transcription profiles. A) Neutrophil clusters that dramatically proportional changed during COVID19 progression. B) Neutrophil clusters that significantly (adjusted P-value <0.05) associated with the severity of COVID19. Spearman’s’ correlations were used to determine the association between cluster proportion and the severity of COCID19 (healthy 1, mild 2, severe 3, decreased 4). Red: positive correlation; Blue: negative correlation; Depth of color increase with the absolute value of the association. Only significant associations (adjusted P-value <0.05) are shown in this graph. C) GSEA analysis of significantly dysregulated genes in selected neutrophil clusters from COVID19 patients compared with that from healthy controls. D) GSEA analysis of significantly dysregulated genes in selected neutrophil clusters from severe COVID19 patients compared with that from mild COVID19 patients.

Collectively, our data demonstrate that distinct clusters of mature neutrophils exist even under basal uninfected conditions, as reflected by distinct transcriptional profiles and activated pathways. The relative proportions of each cluster change during infection and with increasing severity. Furthermore, neutrophils display dynamic responses during infection, with evidence of increased oxidative stress and ribosomal pathway activation during severe infections and suppression of multiple viral pathways (e.g., influenza and measles). Our findings contradict the notion that neutrophils are a homogenous population of transcriptionally limited effector cells and suggest that further investigation into identifying the factors that regulate the development and activation states of different neutrophil subtypes is warranted. Ongoing studies are being conducted to analyze the mechanisms by which different neutrophil subsets emerge, and how each cluster contributes to disruption of epithelial cell barrier function, viral clearance, and resolution of inflammation.

## Methods

### Patient cohort, biological samples, and preparation of single-cell suspensions

Total 11 COVID-19 patients and 5 healthy control in this study were enrolled from the Ann Arbor VA Healthcare System and Michigan Medicine. The 11 patients with COVID-19 were classified into two groups based upon severity – “mild” (n□=□4, hospitalized but needing <50% O2), or “severe” (n□=□7, hospitalized but needing > 50% O2 or in Intensive Care Unit). The demographic and disease characteristics of the prospectively recruited patients studied by scRNA-seq are listed in Supplementary figure 1. All participants provided written informed consent for sample collection and subsequent analyses.

Blood was processed immediately after collection. For PMBC and granulocyte isolation from patient blood, 5ml of blood was carefully layered on 5ml of Lympholyte-poly isolation media (NC9950836, Fisher) in a 15ml conical tube in a biosafety cabinet. The sample was spun in a sealed bucket at room temperature, 500g, 35 minutes. After centrifuge, leukocyte bands containing mononuclear cells and granulocytes were transferred to another 50 ml conical tube. Cells were diluted with an equal volume of HBSS without Calcium and Magnesium and spun at room temperature, 350g for 10min. The supernatant was removed from each pellet and suspended in 5ml 1x ACK lysis buffer for 2 minutes on ice. Pellets were suspended with PBS containing 0.04% BSA and then counted. An aliquot of mononuclear and granulocyte cells was processed in a cytospin and stained using Diff Quick (26096-25, Electron Microscopy Sciences). Cell differentials were counted, and the mononuclear and granulocyte populations were combined to achieve a 1:1 ratio. Cells were immediately processed for the single-cell RNA library. For all samples, cell viability exceeded 90%.

### Single-cell RNA library preparation and sequencing

The scRNA-seq libraries were constructed using a Chromium Next GEM Single Cell 3□ Reagent Kit v3.1 (10X Genomics) according to the manufacturer’s introduction. In brief, the cell suspension (700-1,200 cells per ul) was loaded onto a Chromium single cell controller to generate single-cell gel beads in the emulsion (GEMs). Following this, scRNA-seq libraries were constructed according to the manufacturer’s introduction. The libraries were sequenced using an Illumina Novaseq sequencer at Advanced Genomics Core of the University of Michigan using the suggested cycling from 10X Genomics.

### Single-cell RNA-seq data processing

We aligned single-cell RNA sequencing data against the GRCh38 human reference genome and preprocessed using cellranger pipeline (version 6.0.0). A preliminary single-cell gene expression matrix was then exported from cellranger for further analysis. Quality control was applied to cells based on three metrics: the total UMI counts, number of detected genes, and proportion of mitochondrial gene counts per cell. Specifically, cells with less than 500 UMI counts and 200 detected genes were filtered out, as well as cells with more than 20% mitochondrial gene counts. Thereafter, we applied DoubletFinder, which identifies doublets formed from transcriptionally distinct cells (31), to remove potential doublets. The expected doublet rate was set to be 0.075, and cells predicted to be doublets were filtered out. After quality control, a total of 108,597 cells were collected for further analysis.

### Clustering and cell-type annotation

We used Seurat (23) to integrate and cluster the collected single cells from COVID19 patients and healthy controls. The gene counts for each cells were normalized by LogNormalize method, which divides gene counts by the total counts for that cell and multiplied by the scale.factor. The normalized gene counts were then natural-log transformed using log1p function. The top 2000 most variable genes were selected using FindVariableFeatures functions for the clustering of single cells. We used dimensions of reduction 30 and resolution 0.3 for the cluster analysis. We used SingleR ((24)) and human primary cell atlas reference (32) to annotate cell types of single cells. The cell type of the cluster was determined by the dominant cell type in each cluster. The proportion of B cells, T cells, and NK cells in every sample were further calculated to determine the consistency between our data and observed in other published data (33, 34) in the cell type composition difference between severe COVID19 patients and healthy controls.

### Cellular crosstalk analysis

We used iTALK (35) to identify and visualize the possible cellular crosstalk mediated by up-regulated ligand-receptor pairs between each cell type in COVID19 patients. We used the cytokine/chemokine category in the ligand-receptor database for this analysis. We used Wilcoxon rank-sum test to identify the significantly up-regulated genes ((adjusted P-value <0.05 & average log fold change >0.1) for every cell type in the severe and mild COVID19 patients, respectively, compared with healthy controls at day 0. We also identified up-regulated genes ((adjusted P-value <0.05 & average log fold change >0.1) between day 5 and day 0 in recovered mild COVID19 patients and deceased severe COVID19 patients, respectively. We then matched and paired the up-regulated genes against the ligand-receptor database to construct a putative cell-cell communication network using iTALK. The iTALK defines an interaction score using the log fold change of ligand and receptor to rank these interactions.

### Feature genes and pathways for neutrophils

We collected neutrophil cells from mild and severe COVID19 patients at day 0, as well as healthy controls to identify significantly up-regulated feature genes (adjusted P-value <0.05 & average log fold change >0.1) by comparing one type (healthy, mild, or severe) of neutrophil to the rest of neutrophils. The overlap among feature genes identified in healthy, mild, and severe groups was presented in Venn plot using the Venn package. The top 10 feature genes for each type of neutrophil were presented in a bubble plot using the DotPlot function from the Seurat package. These feature genes were then mapped to human protein-protein interactions (PPIs) downloaded from the BioGRID database (version 4.4.197) using R. The KEGG pathways significantly enriched (adjusted P-value <0.05) in feature genes that connected by PPIs were identified using enrichKEGG function from clusterProfiler package (36). The bipartite plot of significant pathways, genes, and PPIs were presented using the Cytoscape tool (37).

### Neutrophil cluster analysis

We integrated neutrophils from mild and severe COVID19 patients, as well as healthy controls to identify neutrophil clusters using resolution 0.5 in Seurat. The significantly up-regulated ((adjusted P-value <0.05 & average log fold change >0.1) feature genes for each neutrophil cluster were identified by comparing one cluster to all other clusters. The top 5 feature genes for each cluster were shown in a bubble plot using DotPlot function from the Seurat package. The KEGG pathways significantly enriched (adjusted P-value <0.05) in feature genes were identified for each cluster using clusterProfiler package. The top 3 significant pathways were shown in the bubble plot using ggplot2 package. The composition changes of each cluster among healthy control, mild and severe COVID19 were identified using student’s t-test and presented using ggpubr package. We define the severity of COVID19 from 1 to 4, in which 1 means healthy, 2 means mild, 3 means severe, and 4 means deceased severe. The association between composition changes of each cluster and severity of COVID19 were identified using spearman’s correlation and the Hmisc package. The significant associations were shown using corrplot package. We further identified differentially expressed genes (DEGs) between severe and mild, mild and healthy respectively, for each cluster that associated with the severity of COVID19. The KEGG pathways significantly enriched (adjusted P-value <0.05) in DEGs were identified using the gseKEGG function from clusterProfiler package. The normalized enrichment score (NES) of significant pathways indicates the activation status of the pathway.

## Statistics

Statistical analysis was performed using R with Student’s t test or analysis of variance (ANOVA). Asterisks on figures indicate statistical significance as follows: *P < 0.05, **P < 0.01, ***P < 0.001, and ****P < 0.0001.

## Ethics and Research Approval

This study was approved by the VA Ann Arbor Institutional Review Board (IRB) and University of Michigan IRB (IRB-2020-1228 and HUM00181804, respectively). All participants provided written informed consent for sample collection and subsequent analyses. Study procedures adhered to full ethical and safety standards.

## Author Contributions

Experiments were performed and data were analyzed by JX, BH, LXG, and JCD. Experimental support and methods: BH, JX, KC, DV, and JCD. Writing and revision or serious assistance to writing and revision: JX, BH, LXG, MAO, and JCD. Project supervision: LXG, MAO, and JCD.

## Data Availability

the scRNA-Seq data, along with patient severity labels, will be deposited to GEO [in progress].

## Acknowledgments

Funding: LXG is supported by NIH/NIGMS, R01 LM012373 and R01 LM012907 awarded by NLM, and R01 HD084633 awarded by NICHD. MAO is supported by VA ORD RCS 1IK6 BX003615-01 and Merit BX000656Awards, JCD is supported by NIH/NIA R01 AG028082, VA ORD I01BX004565, and I01BX005447.

