## Supplemental Fig 1-5 for "ScRNA-Seq study of neutrophils reveals vast heterogeneity and breadth of inflammatory responses in severe COVID-19 patients"

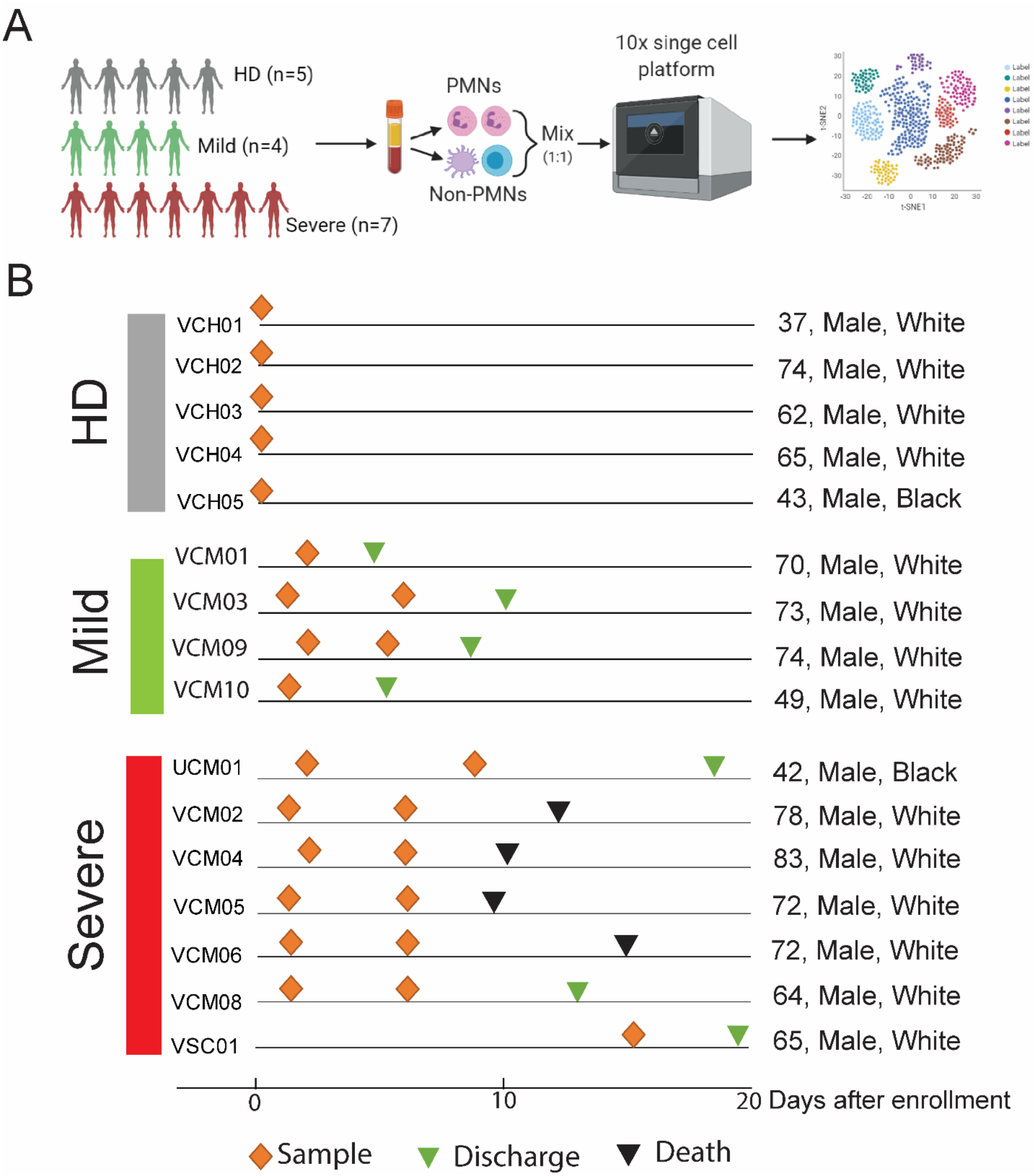


Fig. S1 (A), schematic showing the overall study design. The scRNA-seq was applied to whole blood cells across three conditions and the output data were used for expression analyses. (B), Timeline of the course of disease for 11 patients infected with SARS-CoV-2 enrolled in our study.

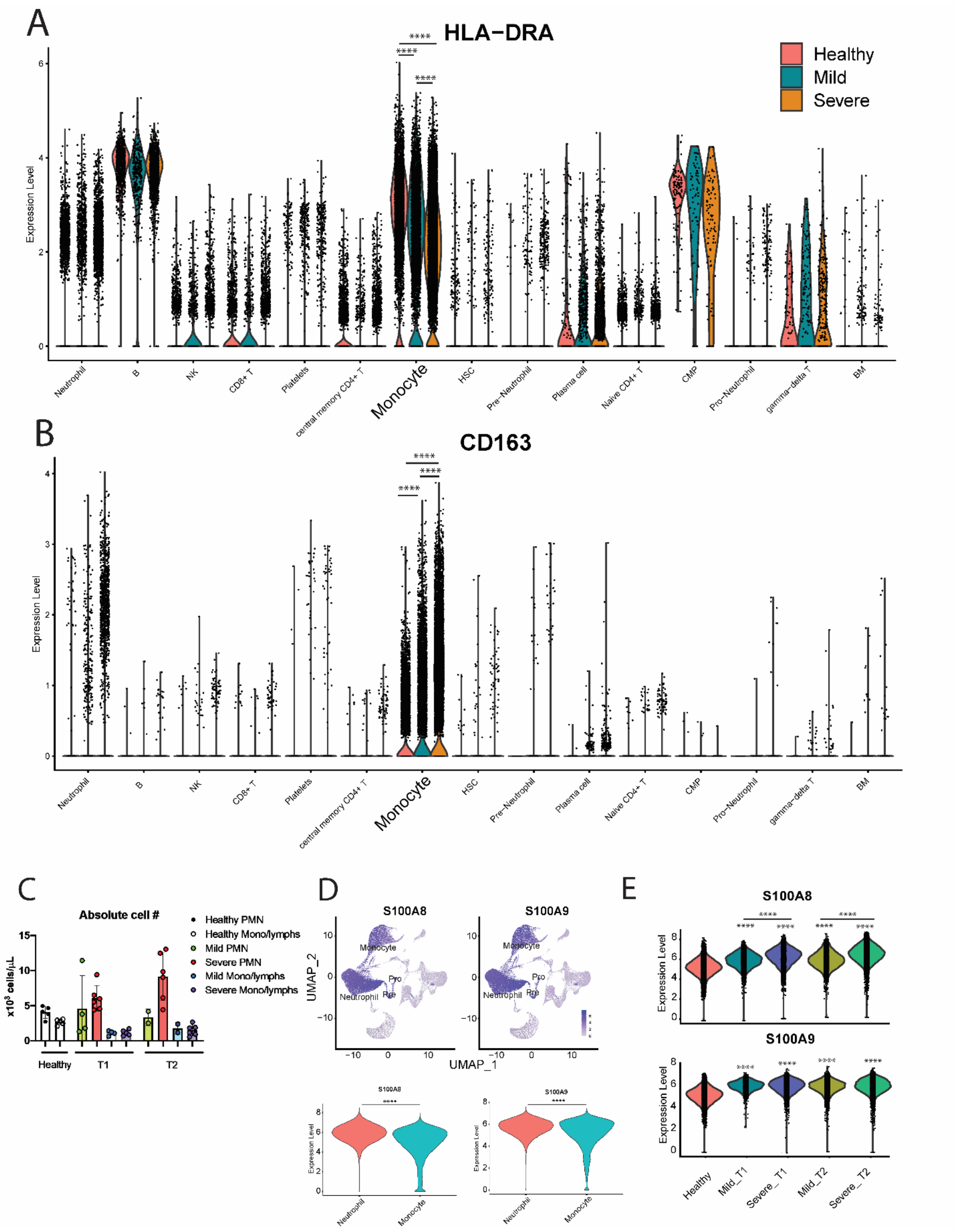


Fig S2. (A) Violin plots showing expression of HLA-DRA by different cell clusters. (B) Violin plots showing the expression of CD163 by different cell clusters. (C) Absolute cell number counts in the blood from healthy, mild, and severe patients (D) UMAP plot showing the expression of S100A8 and S100A9 by neutrophils and monocytes. (E) Violin plots showing expression of S100A8 (up) and S100A9 (below) by neutrophils in different groups of patients. Asterisks on figures indicate statistical significance as follows: ****P < 0.0001 compare to healthy controls.

Fig. S3

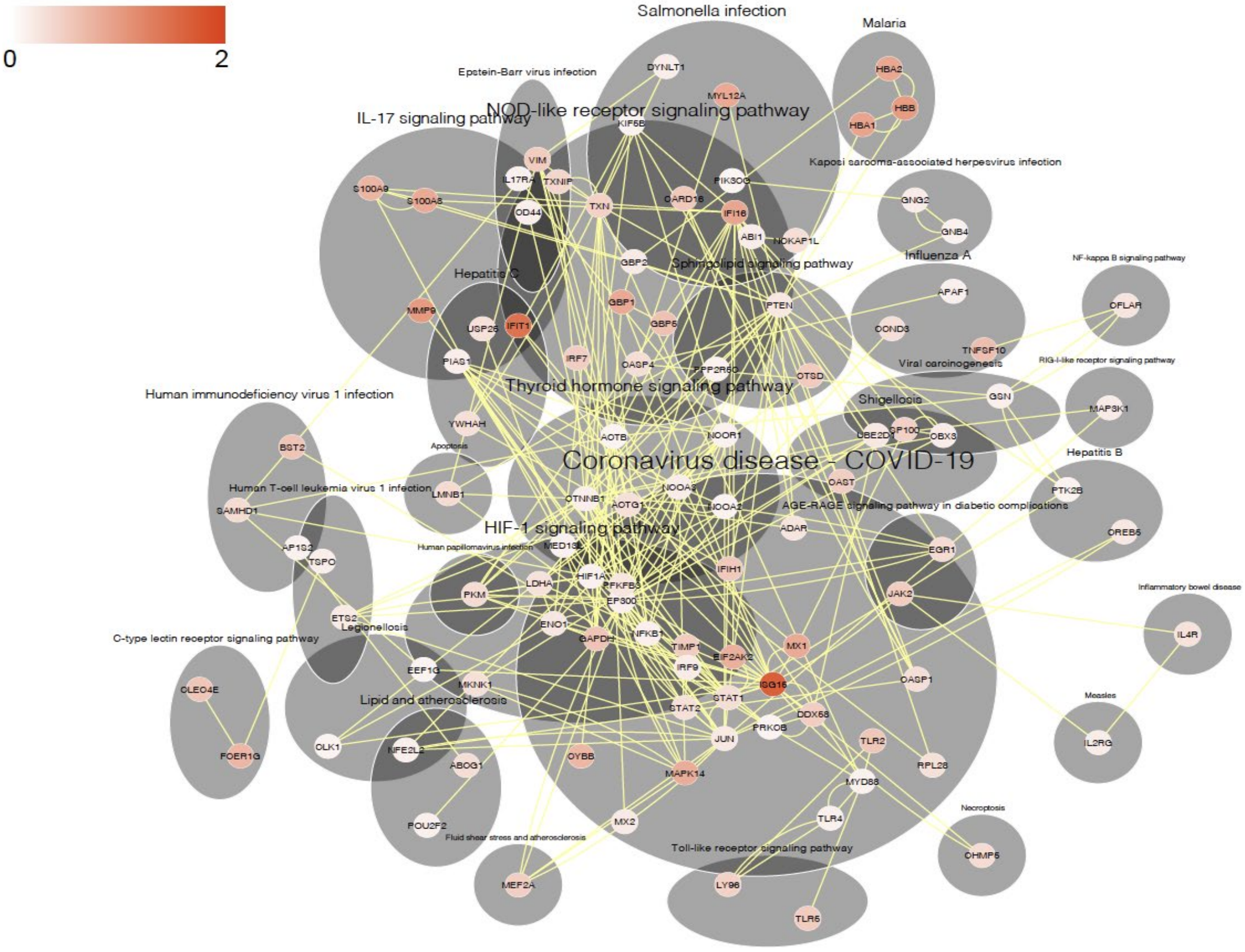


Fig. S3 (A) Protein-protein interaction networks of significantly up-regulated pathways (adjusted P-value <0.05) in neutrophils from COVID19 patients compared with that from a healthy control.

Fig. S4

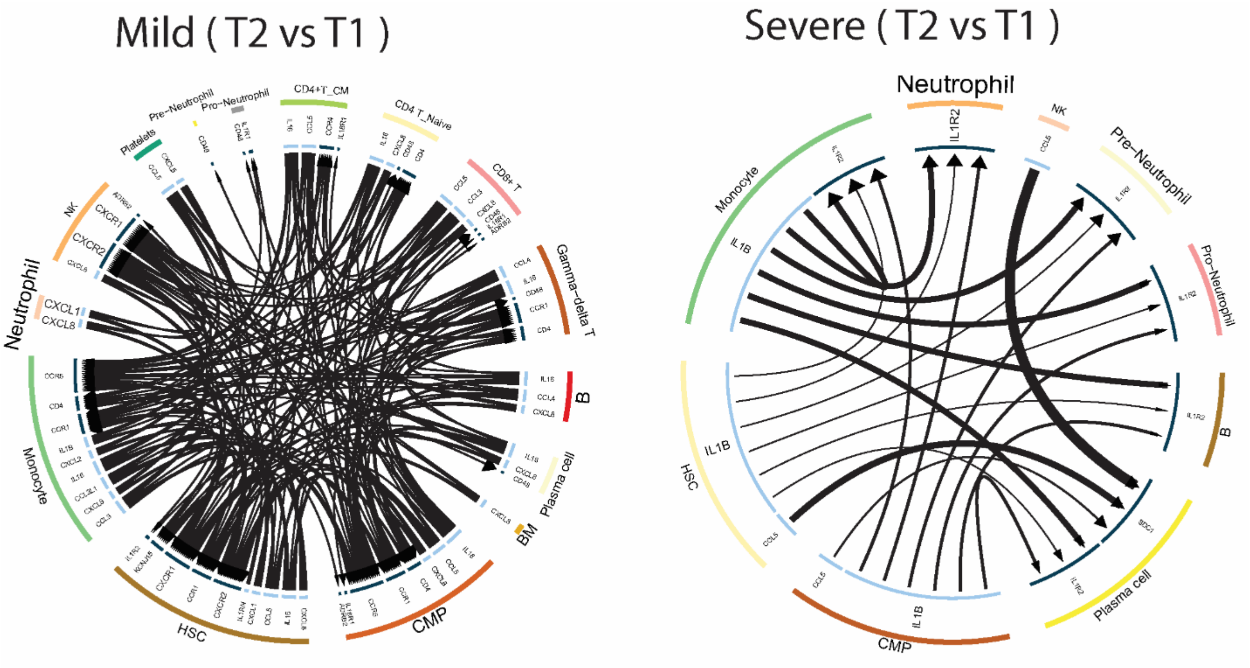


Fig. S4 Circos plot showing the cellular crosstalk mediated by up-regulated (T2 vs T1) ligand-receptor pairs between inflammation-related cell types from recovered mild (left) and deceased severe (right) COVID19 patients, respectively.

Fig S5

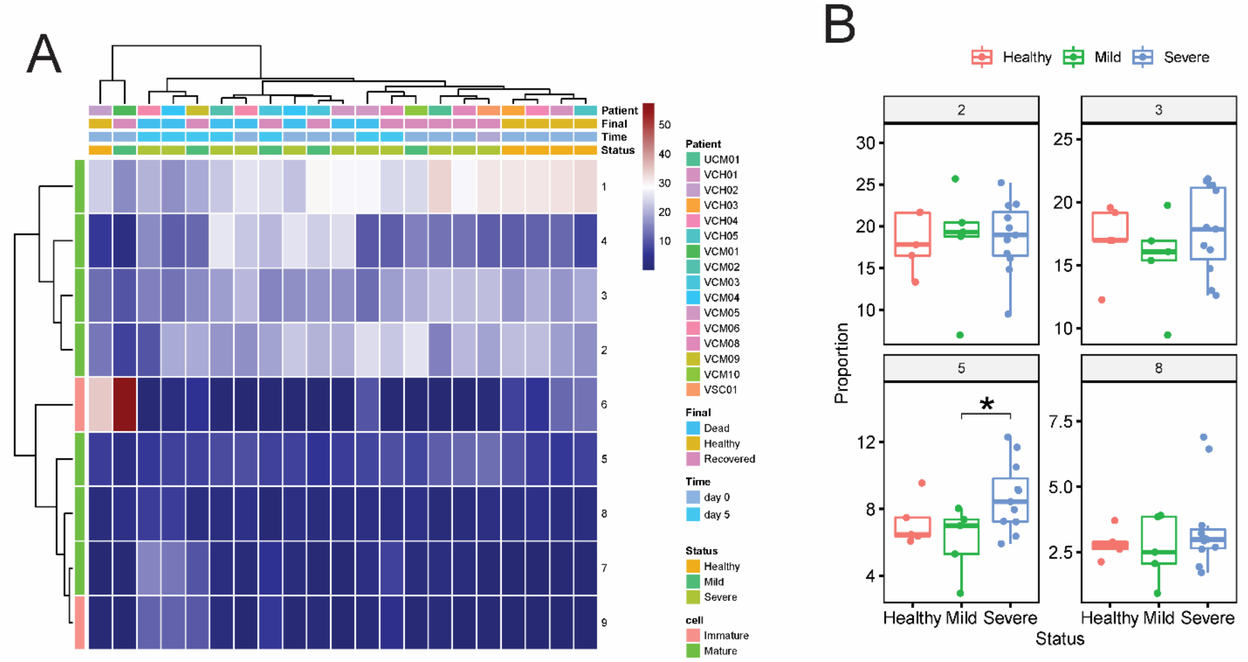


Fig S5 (A) Heatmap of neutrophil cluster size in every sample. (B) Proportional change of neutrophil clusters during COVID19 progression.
